# A Potential Beta-hydroxybutyrate Therapy for Duchenne Muscular Dystrophy in Disease Models

**DOI:** 10.1101/2025.10.18.683257

**Authors:** Sophia Xing-Shan Chen, Jia Lv, Lingfang Tang

## Abstract

Duchenne muscular dystrophy (DMD) is a common and lethal muscular degenerative disorder caused by mutations in the *Dystrophin* gene. Patients with DMD suffer progressive motor and respiratory decline over time and exhibit mitochondria defects at the cellular level, contributing to their reduced life expectancy. Despite extensive efforts, effective treatments for DMD remain unavailable. This study aims to explore a dietary approach to treat DMD, by precisely utilizing ketone bodies, an alternative fuel and signaling molecule for the mitochondria. As the experimental design, beta-hydroxybutyrate (BHB), the ketone body that is most abundant and stable in mammals, was supplemented in the drinking water of DMD disease model mice (*mdx* mice) as well as wild type mice. The results show that exogenous BHB supplementation in *mdx* mice led to significant decreases in plasma creatine kinase (CK) levels compared to chow-fed *mdx* mice, indicating mitigation of muscle damages. Functional tests also show that BHB improved muscle strength and endurance in *mdx* mice, while histological analyses revealed muscle health restoration, evidenced in uniform fiber diameter. BHB supplementation did not adversely affect the growth and overall health in either wild type or *mdx* mice. These findings may provide the proof-of-principle evidence for the use of BHB as a potential dietary approach for DMD treatment.

## Introduction

### Duchenne Muscular Dystrophy

Duchenne muscular dystrophy (DMD) is a severe and inherited muscular degenerative disorder, wherein the patients usually suffer from continuous muscular damage, eventually progressing into severe respiratory and heart problems. It was estimated that this disease has a global prevalence of 4.8 per 100,000 people (Salari et al., 2022). The median life expectancy for DMD patients is 22 years (Broomfield et al., 2021), bringing severe social and economic distress to the patients’ families.

DMD patients suffer muscular weakness, associated motor delays, ambulation loss that leads to wheelchair dependency, respiratory impairment and cardiomyopathy, cumulating into respiratory or heart failure that eventually lead to death (Birnkrant et al., 2018; Wilson, Tinker & Iskratsch, 2022). A biochemical indicator for DMD is the increase in serum creatine kinase (CK) levels. High levels of CK are related to muscle damage and have been commonly reported in DMD patients (Barp et. al., 2021; Birnkrant et al., 2018).

### Pathogenesis

Mutations in the *Dystrophin* gene is the primary cause of DMD. This results in lack of the large structural protein Dystrophin. Dystrophin assembles the dystrophin glycoprotein complex (DGC), providing a strong mechanical link between the extracellular matrix and the intracellular cytoskeleton that reinforces and stabilizes myofibers while playing a role in signal transduction. Dystrophin has an important role in maintaining the integrity of DGC as well as the sarcolemma (Lapidos, Kakkar & McNally, 2004; Wilson, Tinker & Iskratsch, 2022).

The disruption of Dystrophin in DMD patients leads to sarcolemma damage during contraction and causes excess Ca^2+^ flux into the cell, triggering cell death (Wilson, Tinker & Iskratsch, 2022). Consequently, the damaged muscular tissue will develop chronic inflammation and eventually be replaced by fibrotic and fat tissues (Verhaart, & Aartsma-Rus, 2019).

Patients with DMD also exhibit defects in the mitochondria of their muscle cell (Budzinska, Zimna & Kurpisz, 2021). Mitochondria are the cellular organelle responsible for producing and supplying ATP, the energy currency in living organisms. Mitochondria dysfunction is linked to the failure of energy supply and dysregulated cellular homeostasis in DMD. Multi-leveled mitochondria defects are present in DMD, cumulating in the increased secretion of reactive oxygen species (ROS) and heightened oxidative stress (Casati et al., 2024).

### Status-quo of DMD Treatment

To date, there is no effective treatment to cure the disease. Glucocorticoid treatment remains the standard care to manage the symptoms of DMD, associated with the side effects of this hormonotherapy (McDonald et al., 2018; Henricson et al, 2013). Novel therapies have been explored to restore the dystrophin gene, such as supplementing the gene using adeno-associated viruses (AAV), exon skipping using antisense oligonucleotides, stop codon readthrough, genome editing approaches including CRISPR/Cas9, stem cell therapies, and upregulation of dystrophin surrogate proteins. Besides, there are therapies directed to the secondary pathology of DMD, which include compounds for anti-inflammatory treatments, antioxidant treatments, myostatin inhibition, vasodilation, antifibrotic treatments, calcium homeostasis and mitochondrial function. However, concerns regarding the safety, efficacy, applicability and longevity of these therapies remain (Verhaart, & Aartsma-Rus, 2019; Markati et al., 2022; Zhang & Wu, 2019; Shi, Li & Ma, 2023).

In 2023, FDA approved the world’s first recombinant gene therapy for treating certain DMD patients. The drug Elevidys delivers into the body cells a gene that transcribes and translates micro-dystrophin, which is a shortened protein that comprises of selected domains of the normal Dystrophin. Nevertheless, the treatment could only bring clinical benefit to young patients, and its use was thus limited to DMD patients aged 4 to 5 years (Gu, Li & Xi, 2024; Office of the Commissioner, 2023).

### Studies Regarding Ketone Bodies

Of note, ketones are receiving attention for their muscle-promoting roles (Khouri, Ussher & Aguer, 2023). Ketone bodies are primarily produced from fats in the liver and transported to fuel extrahepatic tissues, in the state of fasting, long-period exercise or low carbohydrate availability. There are three types of ketone bodies: beta-hydroxybutyrate (BHB), acetone and acetoacetate (AcAc). They serve as the alternative energy source for various organs, including the heart, brain and the skeletal muscle (Puchalska & Crawford, 2017; Newman & Verdin, 2017; Wei et al., 2023).

Ketogenic diet (KD) is characterized by low carbohydrates, moderate protein and high levels of fat. Although developed initially for treating epileptic seizure, it has been adopted by certain population for weight control and muscle building. While numerous studies have reported the benefits of KD, the changes were generally not statistically significant after 12 months of therapy. The diet’s sustainability and long-term risks still need further investigation (Batch et al., 2020).

Scientists have been increasingly exploring ketones’ potentials in DMD treatment. Fujikura et al. (2021) conducted a study using a ketogenic diet with medium-chain triglycerides (MCT-KD) on rat models of DMD. They discovered that the MCT-KD led to a significant increase in muscle strength as well as diameter of myofibers in the DMD rat model. The MCT-KD also significantly suppressed key symptoms of muscular dystrophy, from muscle necrosis and inflammation to fibrosis. Besides, the diet stimulated the proliferation of muscle satellite cells, an indicator of enhanced muscle regeneration. The impact of MCT-KD on DMD progression was long-lasting for up to nine months.

In another study, Zou et al. (2016) investigated the non-metabolic role of AcAc in promoting proliferation in muscle cells and regulating muscle cell function. They found that AcAc was capable of accelerating muscle regeneration in normal mice through activation of a specific signaling pathway, and it mitigated muscular dystrophy in *mdx* mice.

Parker et al. (2018) examined the changes in skeletal muscle cell physiology and mitochondrial function after the treatment of BHB. Overall, BHB induced beneficial changes for murine myotube mitochondria, namely, enhanced muscle viability, an increase in mitochondrial respiration, no loss of ATP production, and reduced H_2_O_2_ emission. Besides, the addition of BHB resulted in more mitochondrial fusion, which may have occurred because of decreased ceramides.

### Research Gap

In summary, extensive efforts have been devoted to the development of DMD therapy, ranging from dietary, hormonal, to gene therapies, but so far fail to generate an effective therapy. While ketogenic diets have yielded positive effects in one study, DMD patients often exhibit metabolic dysregulation that may be exposed to additional complications from the high levels of fat and proteins in ketogenic diets. Accordingly, precise administration of ketones directly may offer a safe and effective strategy to leverage the potential therapeutic benefits of ketones. Such approach may also shed lights on developing effective treatment to other neuromuscular diseases, or even to mitochondria-related diseases in general. To this end, we chose to examine BHB, the most abundant and most stable type of ketone bodies in mammals.

This study aims to investigate the potential of BHB to mitigate muscular dystrophy in DMD mouse models. We divided wild type mice and DMD disease model mice (*mdx* mice) into the chow diet group and the group that received BHB supplementation. The effect on mitigating muscle damage was monitored by the plasma CK levels and eventually evaluated by behavioral tests and histological examination.

## Materials and Methods

### Animal Model

*Mdx* mice purchased from Cyagen were used as the DMD animal model in this study. The mice have a C3197T point mutation in the *Dystrophin* gene in the C57BL/6J mouse line, which generates a premature stop codon and leads to the expression of truncated non-functional dystrophin. The *mdx* mice are viable but develop muscle dystrophy.

### Experimental Design

C57BL/6J wild type (WT) mice and *mdx* mice were randomly divided into the following groups, with 8 animals in each group: WT-chow, WT-BHB, mdx-chow, and mdx-BHB. Starting from the age of three months, the two BHB groups received supplementation of BHB in the drinking water, with the ratio of BHB amount per day to body weight being 9 g/kg. The two chow groups serve as the control of no treatment.

The BHB supplementation lasted from Week 1 to Week 20, and was continued from Week 24 to 25 after a brief interval.

### Data Collection

Body weight was measured every week and the health status of mice was regularly monitored. At Week 0, 2, 4, 7, 10, 13 and 16, blood was taken from tail veins for the measurement of plasma CK level, using the creatine kinase test kit produced by Zhongsheng Beikong Biotechnology Company. At Week 4, 7, 10, 13 and 16, blood was also taken to measure the plasma BHB level, using the Solarbio BC5060 kit.

All mice were then subject to functional tests, including the treadmill test and the grip strength test. The mice were trained to get familiar with the tests beforehand and performed the tests at Week 17 and 18.

The treadmill was set with an initial speed of 15 m/min and acceleration of 10 m/min. The testing session spanned for 45 min, and the maximum time that each mouse could remain running was recorded, in seconds. For the grip test, each mouse was tested five times for four-limb grip strength, and the peak force was recorded.

At Week 26, the mice were sacrificed. Tibialis anterior (TA) muscles were obtained and weighed before hematoxylin and eosin (HE) staining. Muscle cross section area was analyzed using the software ImageJ.

### Statistical Analysis

Without special mention, all collected data were analyzed using unpaired t-tests. Data are represented as mean + SE_M_.

## Results

To assess the general impact of BHB supplementation on growth and health of the experimental mice, we monitored their body weight over time. Figure 1 shows that all four groups of mice exhibit similar body weight increases over time. Chow-fed mice appeared to be slightly heavier than mice receiving BHB supplementation, but the differences were not statistically significant. Moreover, overall health status of the mice receiving BHB supplementation was also comparable to their chow-fed counterparts.

**Figure 1.**
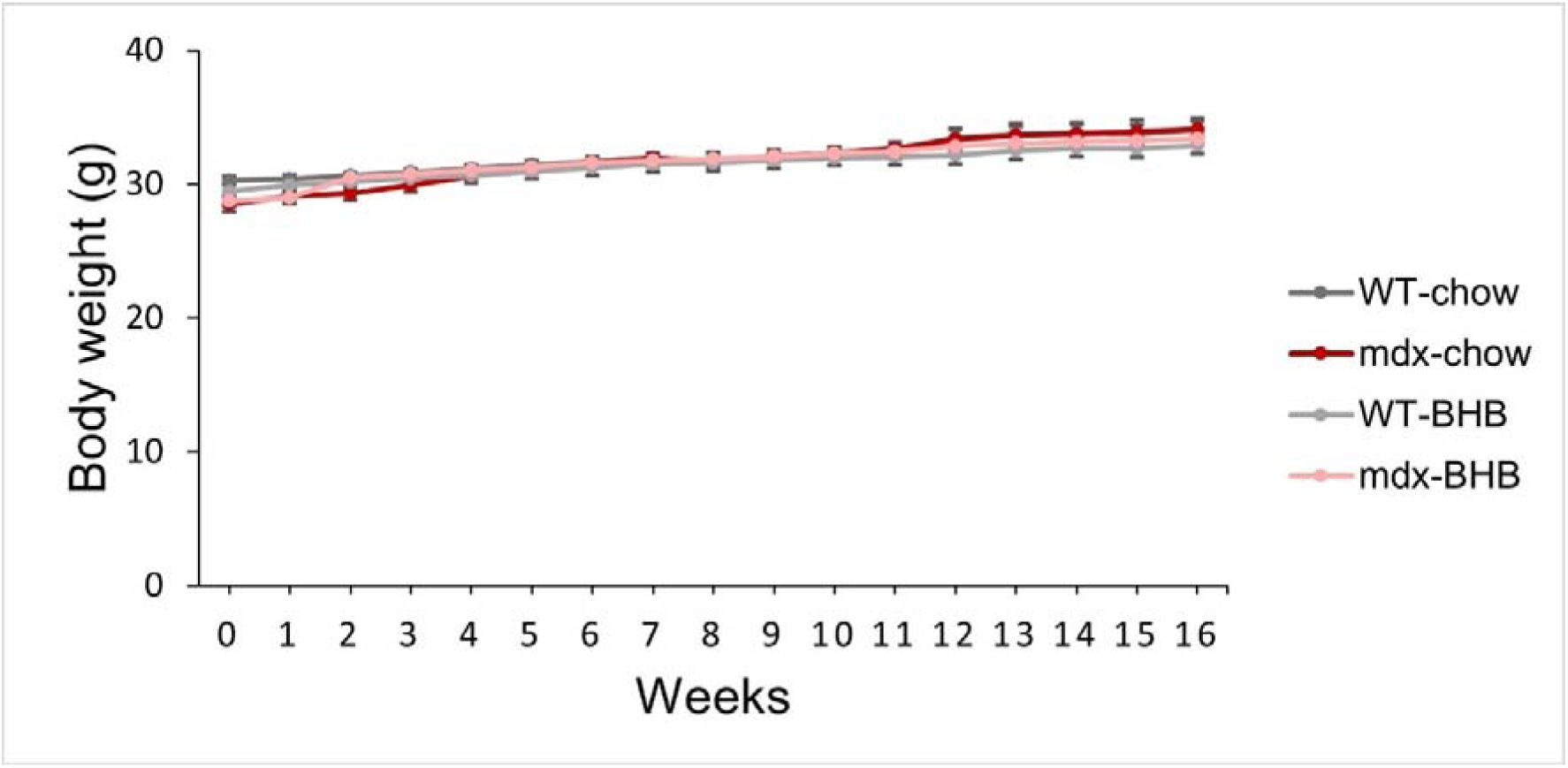
Body weight of mice by week.

To confirm the effect of the dietary supplementation, we monitored plasma BHB levels of the mice every three weeks. Figure 2 shows that BHB levels of the two chow groups remain stable at about 0.5 mmol/L. The two BHB-supplemented groups exhibited a progressive increase of plasma BHB levels, reaching a steady level at about 1.8 mmol/L from week 10 to week 16. Hence, we established the BHB supplementation regimen for the following studies.

**Figure 2.**
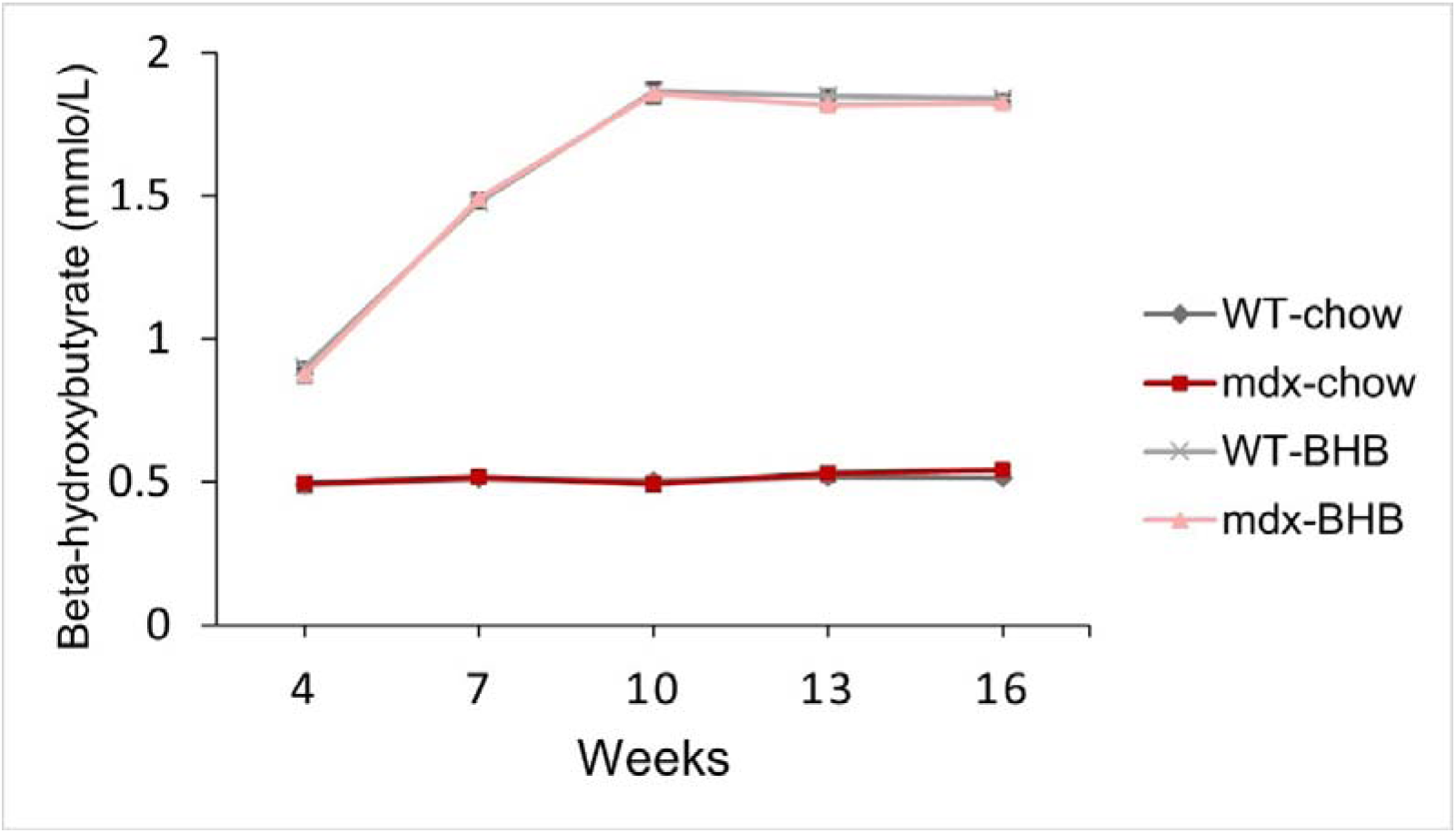
Plasma BHB level of mice at Week 4, 7, 10, 13 and 16.

Using the above regimens, we monitored plasma CK levels to assess the degree of muscle damages. Compared to the wild type mice, *mdx* mice exhibited a marked elevation in baseline levels of plasma CK. At the beginning of the study when the *mdx* mice were three months old, their plasma CK exceeded 20000 U/L, indicating severe muscle damages. The mdx-chow group has its plasma CK level steadily increase over time to over 30000 U/L.

By contrast, the mdx-BHB group initially showed little increase in plasma CK levels till Week 10. Afterwards, the plasma CK levels rapidly dropped to 5400 U/L and decreased slightly thereafter (See Figure 3). As high concentration of plasma CK is an indicator of muscle damage, the results show that supplementing BHB in the drinking water effectively alleviated muscle damages in *mdx* mice.

**Figure 3.**
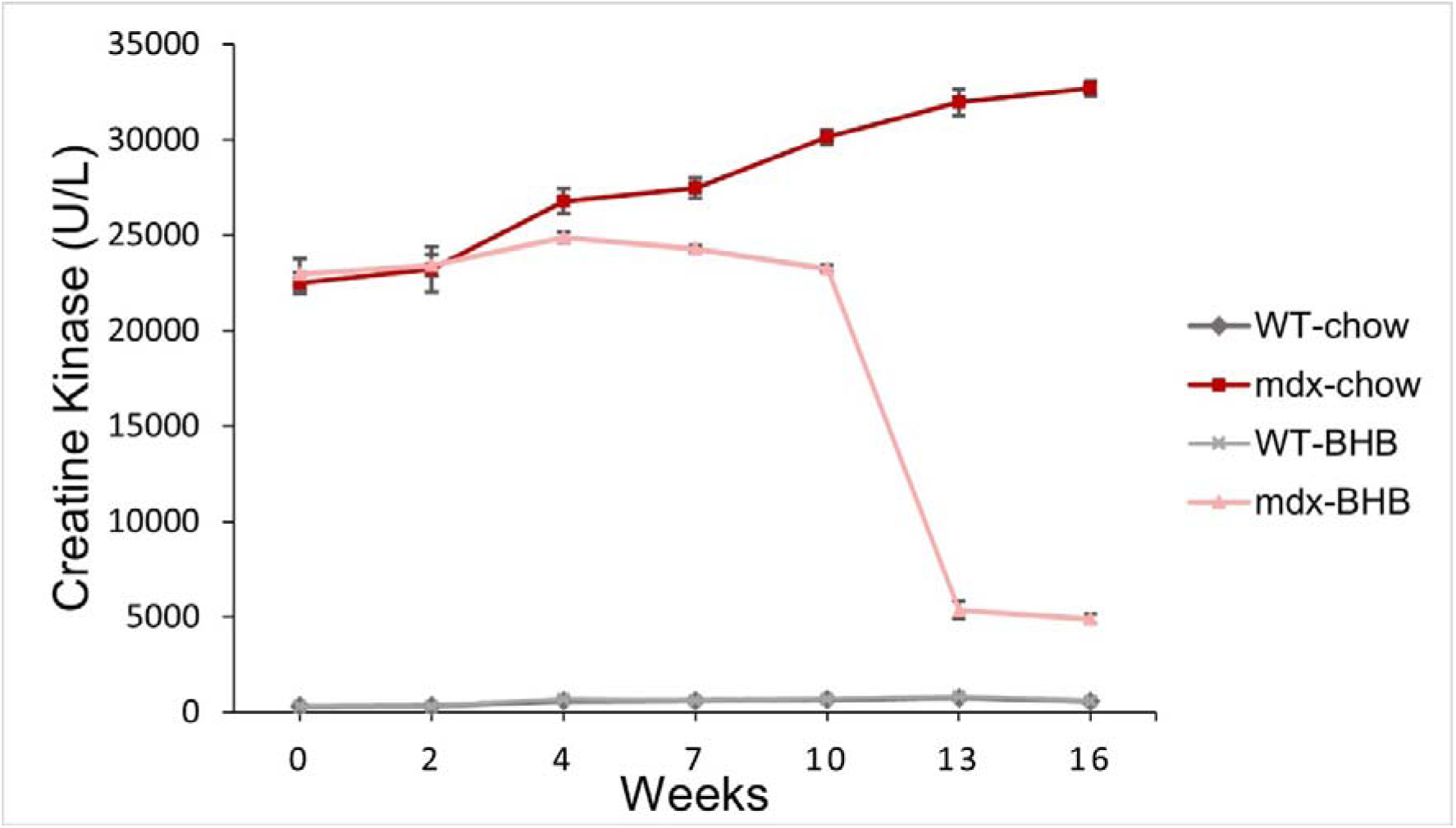
Plasma CK level of mice at Week 0, 2, 4, 7, 10, 13 and 16.

By comparison, the plasma CK levels of wild type mice remain very low throughout the study in both groups, showing that BHB supplementation did not adversely affect the muscle health in wild type mice.

To further evaluate muscle function, we performed functional tests at Week 17 and 18. Figure 4 shows the results of the treadmill tests. Most mice of both wild type groups were able to complete the 45-minute (2700s) treadmill session, regardless of the diet they received. The average maximum running time for mdx-chow group was significantly lower than wild type groups at ∼767s. Importantly, the mdx-BHB group ran nearly twice as long (∼1519s). Thus, the results indicate that BHB supplementation improved stamina in *mdx* mice, although their performance remained below that of wild-type mice.

**Figure 4.**
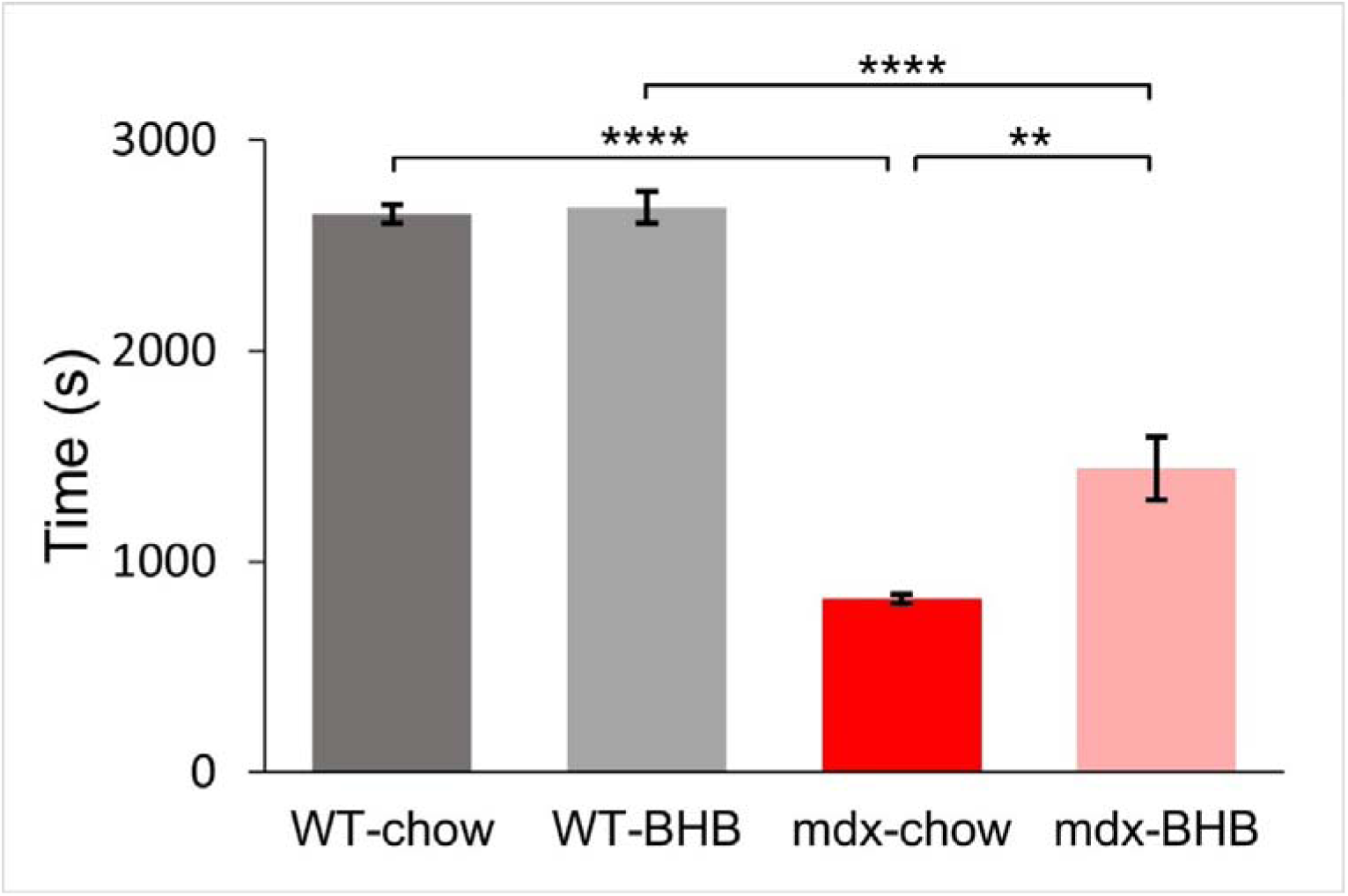
Average maximum running time of mice. **p < .01. ****p < .0001.

Results from the grip strength tests further confirmed the results from the treadmill test (See Figure 5). Both wild type groups have a greater four-limb grip force than mdx groups. However, the mdx-BHB group have a greater four-limb grip force than the mdx-chow group. This result suggests that BHB supplementation improved the muscle strength in *mdx* mice, while it had no impact on wild type mice, which in general showed better muscle health than *mdx* mice.

**Figure 5.**
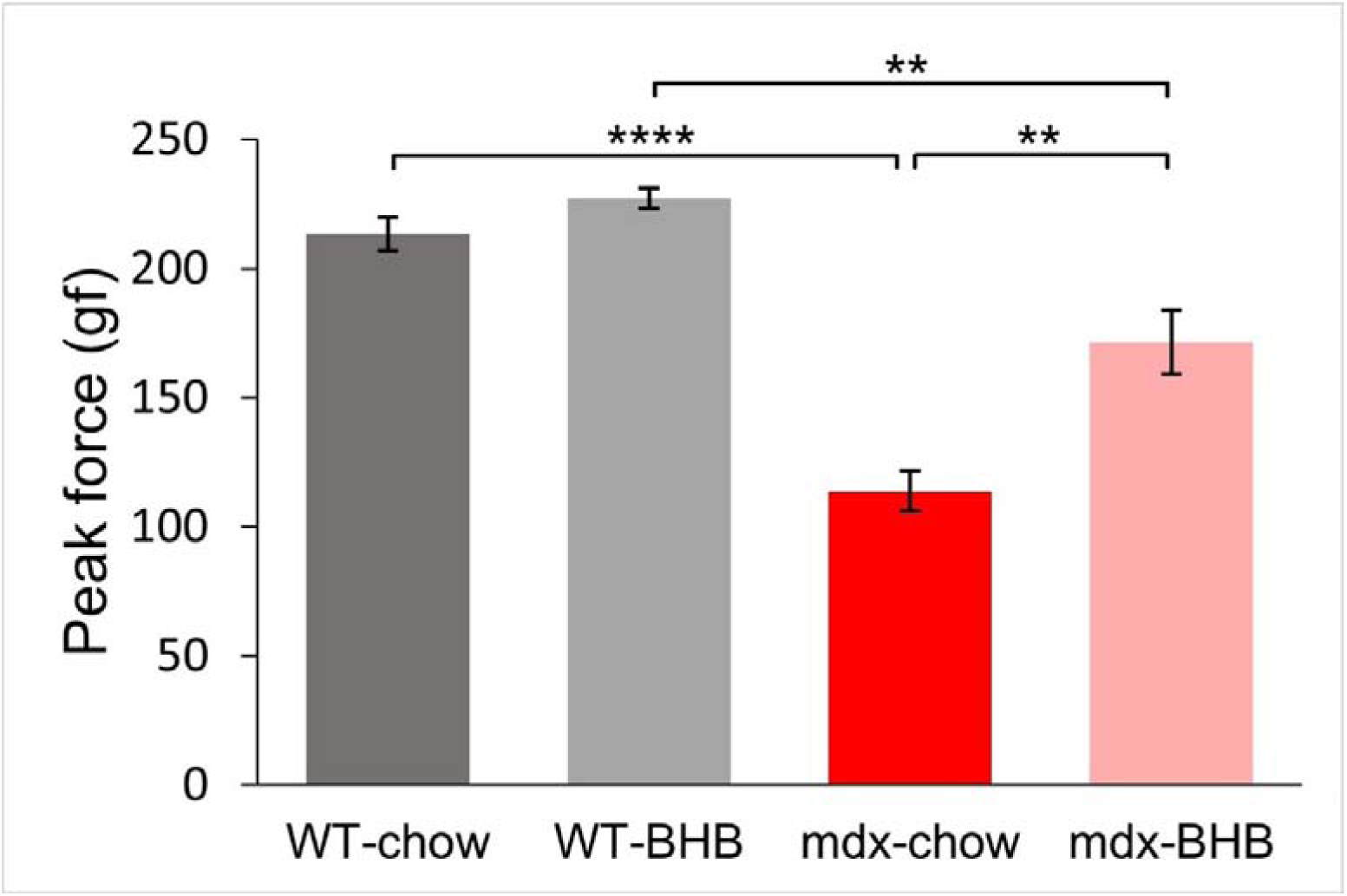
Peak force of mice four-limb grip strength. **p < .01. ****p < .0001.

**Figure 6.**
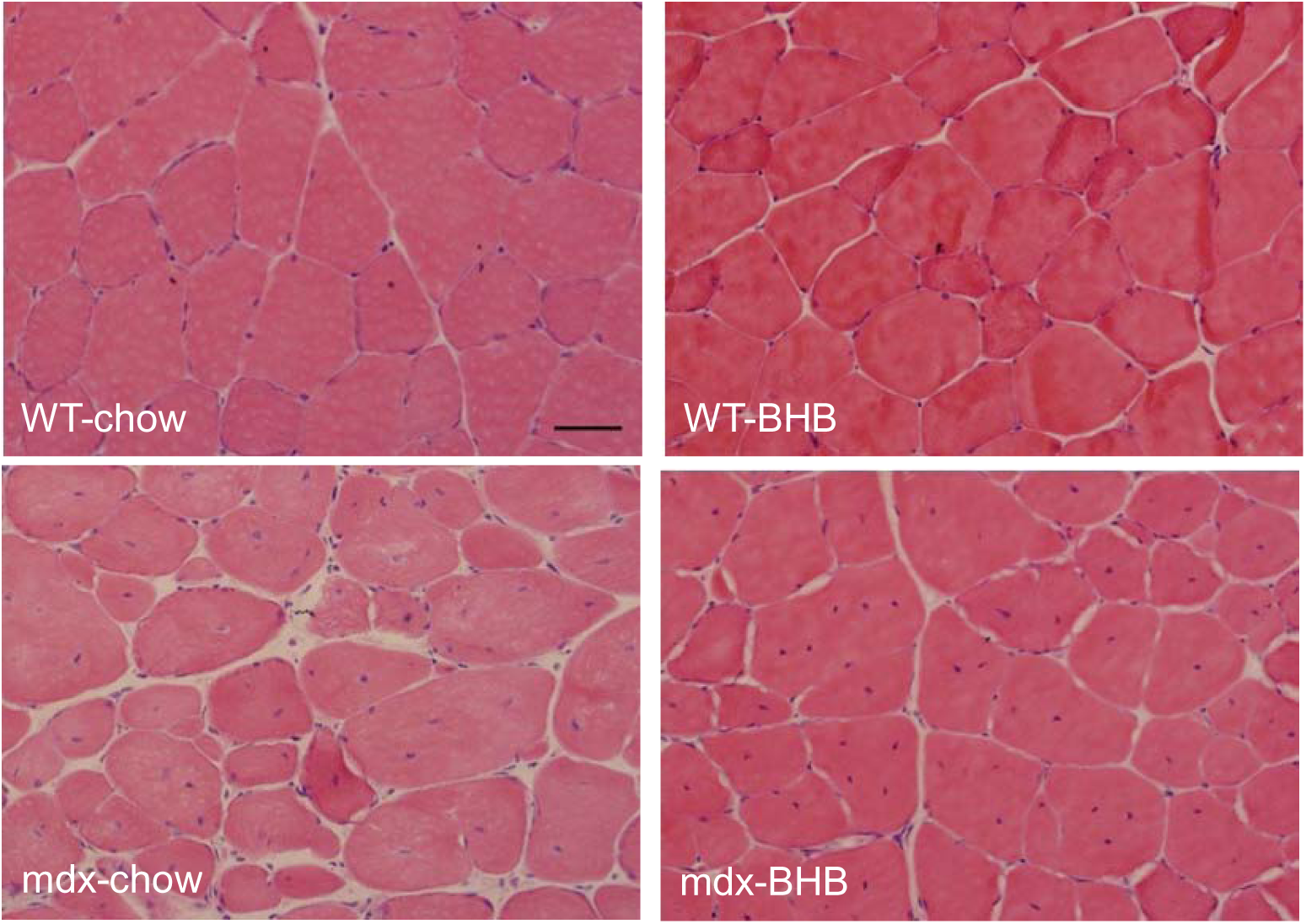
Representative HE-stained TA muscle sections of mice. Scale bar = 200.

**Table 1.** The variances and F-statistics of two-sample F-test for variances.

| Groups | Variance | WT-chow | WT-BHB | mdx-chow | mdx-BHB |
| --- | --- | --- | --- | --- | --- |
| WT-chow | 8252.7 | — |  |  |  |
| WT-BHB | 9769.2 | 0.85 | — |  |  |
| mdx-chow | 32406.2 | 3.93**** | 3.32**** | — |  |
| mdx-BHB | 19175.7 | 2.32**** | 1.96**** | 1.69** | — |

\*\* $p < .01$ . \*\*\*\* $p < .0001$ .

We lastly examined the histological features of skeletal muscles. The WT-chow and WT-BHB groups had normal muscle architecture with relatively uniform fiber sizes, when quantified with their diameters. By contrast, the mdx-chow group mice exhibited marked variation in their muscle fiber diameters, consistent with their weaker muscle strength. Interestingly, a two-sample F-test for variances revealed that the mdx-BHB group had a significantly more uniform fiber diameter compared to the mdx-chow group, suggesting a partial restoration of muscle structure by BHB supplementation.

## Conclusion and Discussion

Using *mdx* mice as a model for DMD, our current study found that dietary supplementation of BHB improved muscular health and function without causing apparent adverse effects, supporting BHB’s potential as a novel dietary treatment for the disease.

Muscle health was assessed by plasma CK levels and functional assays including treadmill running and the grip strength tests. BHB supplement lowered the elevated CK levels in *mdx* mice, reflecting reduced muscle damages. Accordingly, BHB-treated *mdx* mice also exhibited improved running capacity and grip strength. By contrast, BHB supplementation had little influence to the wild type mice. Histological examination of the muscle tissues confirmed that BHB alleviated, to some extent, the muscular dystrophy and tissue damages in the *mdx* mice. However, additional molecular characterizations are needed to fully assess such restoration effects of BHB.

While above results point to a previously unrecognized therapeutic potential of BHB for DMD, the mechanisms remain to be clarified. BHB may act as an alternative fuel source for the impaired mitochondria, or as a signaling molecule that could promote muscle repair. Whether our study in mice models could be translated into human patients also remains a question, although the simplicity and accessibility of BHB supplementation may facilitate future clinical exploration, with appropriate caution.

